# The coexistence of localized and distributed behavioral information in neural activity

**DOI:** 10.1101/2023.11.17.567603

**Authors:** Gaurang Yadav, Bryan C. Daniels

## Abstract

The degree to which control of an animal’s behavior is localized within particular neurons or distributed over large populations is central to understanding mechanisms of decision-making in brains. A first step in answering this question comes from understanding the scales at which neural activity is predictive of behavior. Here, we demonstrate how information measures at the individual, pairwise, and larger group levels characterize the localization of predictive information. We demonstrate these tools using high-dimensional neural data related to nematode and macaque behavioral decisions. Intriguingly, in both examples we find that similar behavioral information coexists across scales: the same information can be extracted from small groups of individually informative neurons or larger groups of randomly chosen neurons that individually have little predictive power. Our results suggest that methods for causal inference may miss potential causal pathways if they are biased toward finding localized control mechanisms.

## 1 Introduction

It is clear that the behavior of animals is controlled by neural activity. Mechanistic hypotheses for the flow of control from sensory input to behavior are abundant in computational neuroscience, (e.g. [1–6]), and specific neurons or groups of neurons have been shown to be part of this flow of control via causal perturbation experiments in some cases (e.g. [1, 7, 8]). Yet it remains unclear in general whether the antecedents of behavior typically lie at the scale of individual neurons or larger groups. Prior studies in multiple organisms have found information about behavior that can be decoded from small numbers of neurons in the brain [9, 10], and others have found similar information distributed over many neurons [11–13].

Recent advances in obtaining data concurrently from large groups of neurons in behaving animals enable new approaches to this question. Even without perturbations at the neural level that would allow for understanding of causal control, observational data constrain control by giving insight into the scales at which behavioral information is available in neural activity. Here we consider behavioral information as variance in neural activity that is predictive of behavioral states defined at the organismal level, and ask whether this information is localized or distributed in the sense of involving few or many neurons. We also ask whether information may exist redundantly at multiple scales. We focus on behavioral decisions, in which the animal carries out one of multiple pre-existing, distinct possible behaviors, which allows us to more easily use the tools of information theory. We demonstrate the use of these tools on two model organisms for which relevant experimental data already exist.

First, the nematode *C. elegans* stands out as a useful model system due to its simplicity, with a known number of neurons and a largely reproducible network structure [2]. A mechanistic understanding of the neural control of behavior in *C. elegans* has been the goal of a substantial research community in both theoretical and experimental neuroscience for decades [1–3, 11, 14, 15], but a full explanation of the connection between neural activity and movement behavior is still being formulated [1, 15, 16]. The emerging picture is one in which neural activity in a large proportion of neurons, even those thought previously to be more related to sensing, is correlated with behavioral states such as forward and backward crawling [11]. Some individual neurons are more centrally important to locomotive behavior (such as so-called “hub” neurons like AVA) and may act as more centralized locations of control [11, 17]. Yet the entire “motor command” that accounts for body motion is not centralized, but rather distributed over a much larger set of neurons [17]. A recent study measured neural activity from most neurons in the animal’s head during free motion, finding that neural populations could predict behavioral states like velocity and body curvature better than single neurons, with roughly 20 to 40 neurons needed to saturate predictive power [12].

In the much larger primate brain, the neuronal dynamics of decisions leading to a well-defined behavioral output have also been studied for decades. Motion discrimination tasks have revealed areas of the brain that represent motion in the visual field and those that integrate this information to produce a perceptual decision, such as whether dots in a visual field are moving predominantly to the left or right [18–20]. Theoretical interpretations have mostly focused on the distinguishing abilities of individual neurons (that is, the information carried by individual neurons), which is typically characterized by the sensory direction of motion to which a given neuron responds most strongly (in the simplest case also corresponding to the behavioral output direction of a saccade indicating the monkey’s choice). Theoretical understanding of the underlying neural dynamics has focused on models of noisy accumulation of sensory evidence [4–6]. Neurons in relevant areas of the brain can be found that contain information both about incoming sensory evidence and that predict the upcoming decision, with accumulation of sensory evidence over timescales of hundreds of milliseconds. Based on the signal and noise levels found in individual neurons, earlier studies predicted the minimum number of neurons needed to be pooled in order to get to the perceptual accuracy displayed by the monkey, finding that four to eight neurons from area MT would be sufficient [19]. A recent study used neurons sampled from prearcuate gryrus to measure the minimum number of neurons needed to predict behavior, finding dynamics that changed from needing ∼ 20 of the measured neurons before the decision and ∼ 1 neuron near the time of the behavior [5].

An important subtlety in the distributed encoding of information is the extent to which the encoding is decomposable. Information that is available from a group of neurons may be fully decomposable, in the sense that subsets of the group independently carry parts of the group information. At the opposite extreme, the group information may be purely non-decomposable or “synergistic,” such that subsets do not carry any of the group information individually [21, 22]. The importance of synergy in this context is the difference between relevant behavioral information changing gradually with the removal of neurons from an informative group (allowing potentially for more robust control mechanisms) versus a scenario in which predictive power is more severely diminished when removing individual neurons. Measuring the degree of synergy in high-dimensional observational data is challenging. One approach for larger groups of variables, known as the O-information, computes an average degree of synergy or redundancy over an entire system without distinguishing any single variable as the output [23], and has been used to find evidence of synergistic encoding in human [24] and macaque neural data [25]. At the smallest level of pairs of variables, the partial information decomposition can be used to measure the extent to which a particular behavioral output is encoded synergistically by pairs of variables [21].

In this study, we highlight the localization of behavioral decision information as being both straightforward to compute from observational datasets and potentially revealing about neural mechanisms for collective decision-making. Instead of searching for a single or optimal “encoding scheme”, we allow for the possibility that multiple subsets of neurons, of different sizes, may redundantly contain the same information. As an initial examination of the practical utility of this approach, we analyze existing measurements of neural activity in a *C. elegans* nematode during free movement and a macaque monkey during a perceptual decision task. We determine the degree to which behavioral information is localized or distributed across observed neurons, measuring information at the scale of individual neurons, pairs of neurons, and larger sets of neurons. At the pairwise level, “purely” distributed information can be readily measured using an existing definition of synergy, and we explicitly quantify and examine sources of pairwise synergy in each system. At larger scales, we examine the possibility that localized information coexists with distributed information over different subsets of neurons.

## 2 Methods

### 2.1 Study systems and datasets

We use two existing datasets that simultaneously measure behavior and activity over multiple neurons: *C. elegans* during free movement [26] and a macaque during a decision-making task [27, 28].

#### 2.1.1 Nematode dataset

The nematode dataset consists of a recording session for whole-brain recording of calcium activity with cellular resolution in a freely moving *C. elegans* worm. The animal’s neurons expressed the calcium indicator GCaMP6s and the fluorescent protein RFP, which are used to estimate neural activity as a function of time using an existing data processing codebase [29] (https://github.com/leiferlab/PredictionCode). We use recording AML32A of the AML32 strain, with a recording duration of approximately 11 minutes. In this dataset, the worm’s body centerline was tracked and analyzed using eigenworm analysis [30] to automatically classify behavior into three categories: forward crawling, backward crawling, and turning.

#### 2.1.2 Macaque dataset

The macaque dataset consists of recording session V07 from an experiment in which a monkey performs a direction discrimination task with a fixed stimulus duration [27, 28]. The dataset consists of 1206 trials, with 219 simultaneously recorded neuronal units in the prearcuate gyrus area of the prefrontal cortex.

The monkey maintained eye gaze fixation on a central point while viewing random dot motion for 800ms. The motion direction and strength varied randomly per trial. After a variable delay, the fixation point disappeared (Go signal), cueing the monkey to report the perceived motion direction by making a saccade to one of two targets (either right or left). Spikes recorded from each electrode were sorted using standard techniques to identify unique neural units representing the activity of individual neurons or small groups of neurons. Data are publicly available and were downloaded from the Kiani lab website (http://www.cns.nyu.edu/kianilab/Datasets.html). Neural activity was analyzed in a time window spanning 1500 ms before to 1500 ms after the go cue. In some analysis we focus on the time window 300 ms after the go cue, which occurs just after the behavioral output, as the macaques have a mean reaction time near 250 ms [28].

### 2.2 Information measures at the level of individuals and pairs

First, we quantify how much information about the behavior of interest is encoded in each individual neuron by calculating the mutual information between a particular neural unit and the corresponding behavior. Next, for pairs of neural units, we estimate the mutual information between the behavior and the pairwise joint neural activity.

Distinct neural states are defined as follows: For the macaque case, neural spikes are counted in 100 ms time slices, and distinct states are defined using 10 bins equally spaced between 0 and the maximum number of spikes for each neural unit across trials. For the nematode case, distinct states are defined using 5 bins of neural calcium activity levels equally spaced between the minimal and maximal activity level for each neuron across time.

Estimates of mutual information measures rely on estimating the entropy of distributions of measured variables, both individual variables and joint distributions over multiple variables. Estimating the entropy of distributions based on sampled data is challenging given small sample sizes—and even more challenging for this reason when estimating distributions over multiple variables. To get a less biased estimate of entropy than provided by the “naive” estimate that equates measured frequencies with probabilities, we use the NSB method [31], which also provides an estimate of our uncertainty in the entropy. Our code for estimating information measures is available at github.com/Collective-Logic-Lab/InfEst.

In the nematode case, neural calcium levels are measured on a faster timescale than the correlation time of about 5 s [32]. To correctly characterize our uncertainty in the information measures, we subsample the data at 5 s time intervals, producing 137 samples in the time series.

### 2.3 Synergy and redundancy

Information theory techniques involving one and two variables (e.g. entropy and mutual information, respectively) are well understood and widely used. Going beyond two variables to three and more becomes much more complicated, and the best measures are less well agreed upon [21, 23, 33, 34].

In this study, we use the partial information decomposition method for measuring synergy [21, 34]. Here, synergy is defined as the extra information added by knowing two variables together instead of separately. Specifically, the joint mutual information provided by two variables *X*_1_ and *X*_2_ (in our case, neural activity levels) about a third variable *Y* (in our case, a behavioral state) can be decomposed into unique contributions from *X*_1_ and *X*_2_ separately, a redundant contribution common to *X*_1_ and *X*_2_, and a synergistic contribution that is only obtained by measuring *X*_1_ and *X*_2_ simultaneously.

In order to obtain estimates of synergy that are less sensitive to biases due to limited data, here we develop a new simplified synergy measure that can be easily computed using existing methods for estimating the entropy of sampled distributions. In the partial information decomposition, the synergy is defined as the remaining joint information that is not accounted for as unique or redundant [21, 34]:

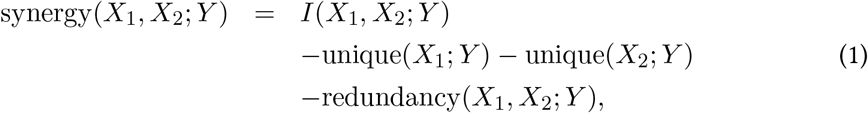

where *X*_1_ and *X*_2_ here represent the activity distributions of two neural units and *Y* is the distribution of behavioral states. Each unique information is itself the difference between an individual mutual information and the redundancy, leading to

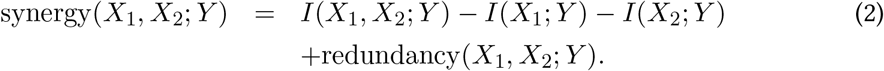

The redundancy is defined in terms of the minimum specific information *I*_spec_(*X*_*i*_; *y*) provided about each behavioral state *y*:

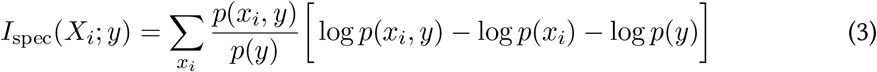

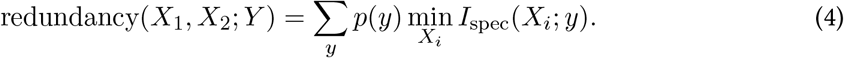

Unfortunately, the specific information cannot be written as a difference of entropies—if it were, then we could use a standard entropy estimation algorithm to estimate the redundancy and therefore the synergy, as all remaining terms are mutual informations that can be expressed as differences in entropies. As a way around this, we note that, if the same *X*_*i*_ were to minimize the specific information across all behavioral states *y*, the redundancy simplifies significantly:

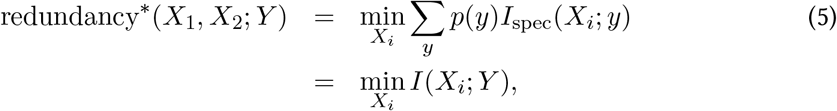

in which case we could use this simplified version of the redundancy to define a corresponding simplified version of the synergy that can be computed fully in terms of individual and joint mutual informations:

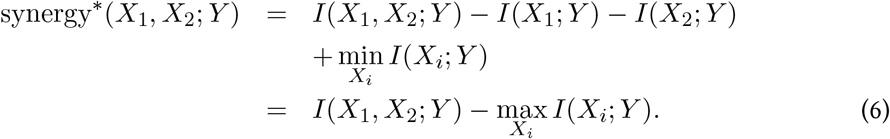

We use this simplified expression to estimate synergies using the NSB method. We can get a rough estimate of the error introduced by this simplification by computing the difference between the original (Eq. 2) and simplified (Eq. 6) synergies using naive estimates of the probabilities of states that simply set them to the frequencies with which they are seen in the data.

### 2.4 Linear discriminant analysis (LDA)

Linear discriminant analysis is a method used in statistics and pattern recognition to find a linear combination of features that separates two or more classes of events [35]. Similarly to principal components analysis, LDA also looks for linear combinations of variables which best explain a given dataset. Instead of finding dimensions of maximum variance, LDA attempts to find the linear combination of variables that best separates two classes of data [36].

Here, following Ref. [5], we use LDA as a computationally tractable method for setting a lower bound on the predictive power contained collectively in the neural states of a set of *N* neurons. In general, if the encoding of behavioral information is nonlinear, LDA will make suboptimal predictions, but it does provide a useful lower bound on the optimal predictive power.

Briefly, we use the most straightforward version of LDA, which outputs a vector 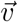 along which neural states can be projected such that the result is maximally correlated with the binary value of the corresponding class. Because the most easily interpreted version of LDA is one that discriminates between two classes, we coarse-grain the behavioral state in the nematode case to two states: forward and non-forward.

To measure predictive performance, we start by splitting data into training and test data, consisting of continuous subsets of time in the nematode case and randomly selected trials in the macaque case. For nematode and macaque both, this process was repeated 4 times. Care was taken to maintain class balance, with the training and test splits containing approximately 88 and 12 percent respectively of the total samples, respectively. The class distribution was approximately equivalent between the train and test splits for each fold. This repeated crossvalidation methodology allowed us to rigorously assess the generalization ability of our approach across multiple train-test splits. When performing LDA for the nematode data, we choose in- and out-of-sample data over contiguous subsets of time. Using this method, correlations over time do not allow for overfitting trial-level noise, so we do not subsample the data as we do when computing the other information measures, instead using all 4083 measured timepoints. After running LDA on given training data, we then test its accuracy on correctly predicting behavioral states on corresponding test data. We use the scikit-learn python package to compute LDA.

## 3 Results

### 3.1 Information in individual neural units

As shown in Figure 1, at the level of individual neurons, behavioral decision information is relatively localized. In both the nematode and macaque examples, there are a handful of neurons whose activity is more informative about the behavior of interest, with most neurons carrying little or no information individually about this behavior. In the macaque case, due to the preponderance of uninformative neurons, we focus on the 100 neural units that have the largest mutual information averaged over trial time. Even for the very coarse behavioral states that we examine here, no single neuron captures all the behavioral information as quantified by the behavioral entropy (1.0 ± 0.1 bits in the nematode example and 0.993 ± 0.004 bits in the macaque example), but multiple individual neurons do represent a significant fraction of this total.

**Figure 1.**
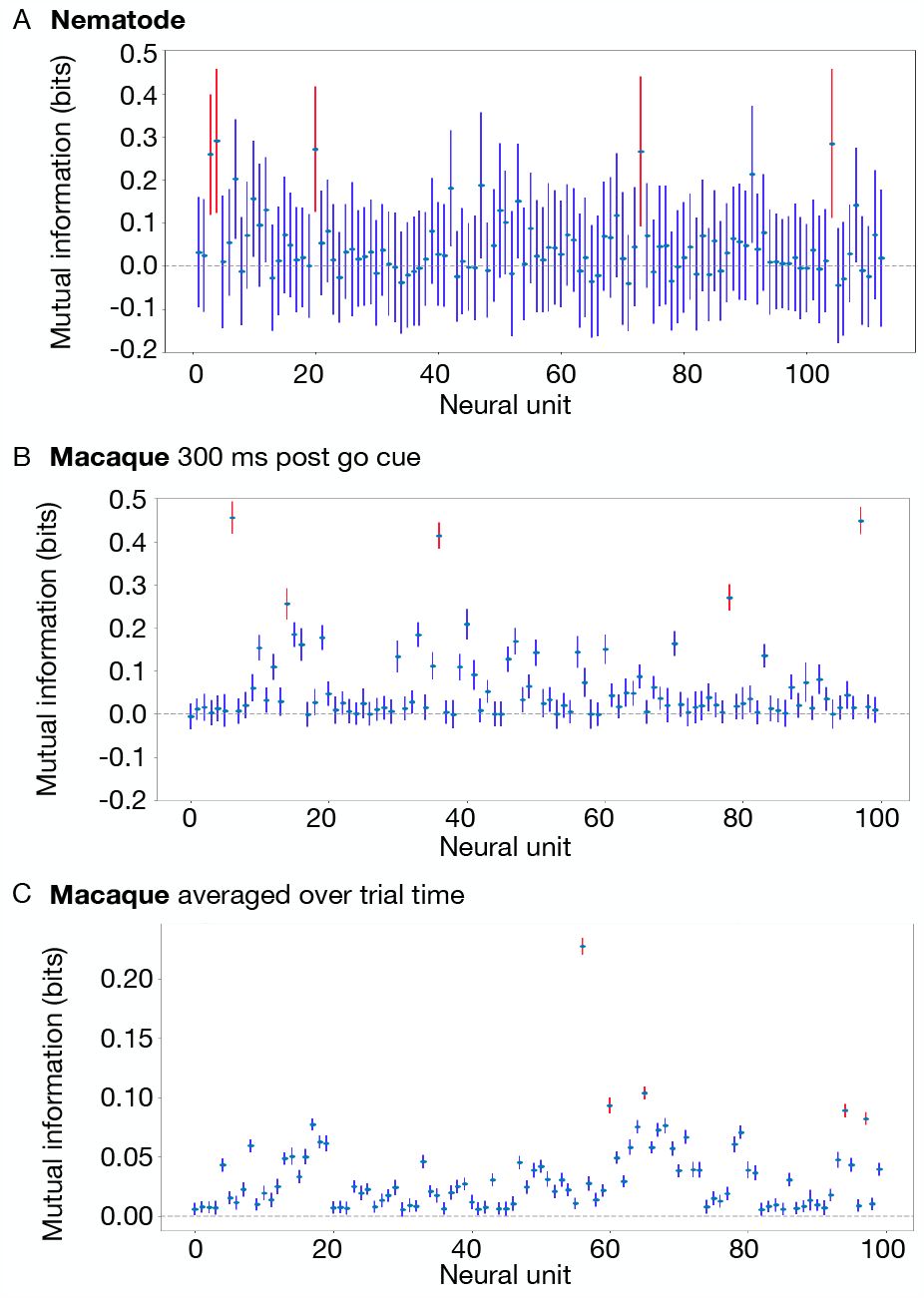
Behavioral information is substantial in particular individual neurons. The mutual information measures the amount of information about behavioral states at the scale of individual neurons, in (A) the nematode, and (B) the macaque near the time of the decision output (300 ms post go cue), and (C) the macaque averaged across trial time. The five individual neural units carrying the most behavioral information in each case are highlighted in red. The mutual information and the uncertainty of the joint entropy (plotted as error bars) are estimated using the NSB method. Note that cases of negative mutual information are consistent with zero under statistical uncertainty.

### 3.2 Information in pairs of neural units: Joint Information, Synergy, and Redundancy

We next examine the ability of pairs of neural units to predict the behavior. We particularly focus on synergistic information, highlighting pairs of neural units that contain more information than can be extracted from each individually. We characterize synergy for all pairs of neural units in Fig. 2. In the nematode example, the limited number of effective data samples increases the statistical uncertainty in our measurement to about 0.2 bits, and we do not find any pairs with synergy significantly above this noise floor. In the macaque example, the larger amount of data gives us more precision, and we do find many pairs with significant synergy, with magnitude of about 0.2 bits.

**Figure 2.**
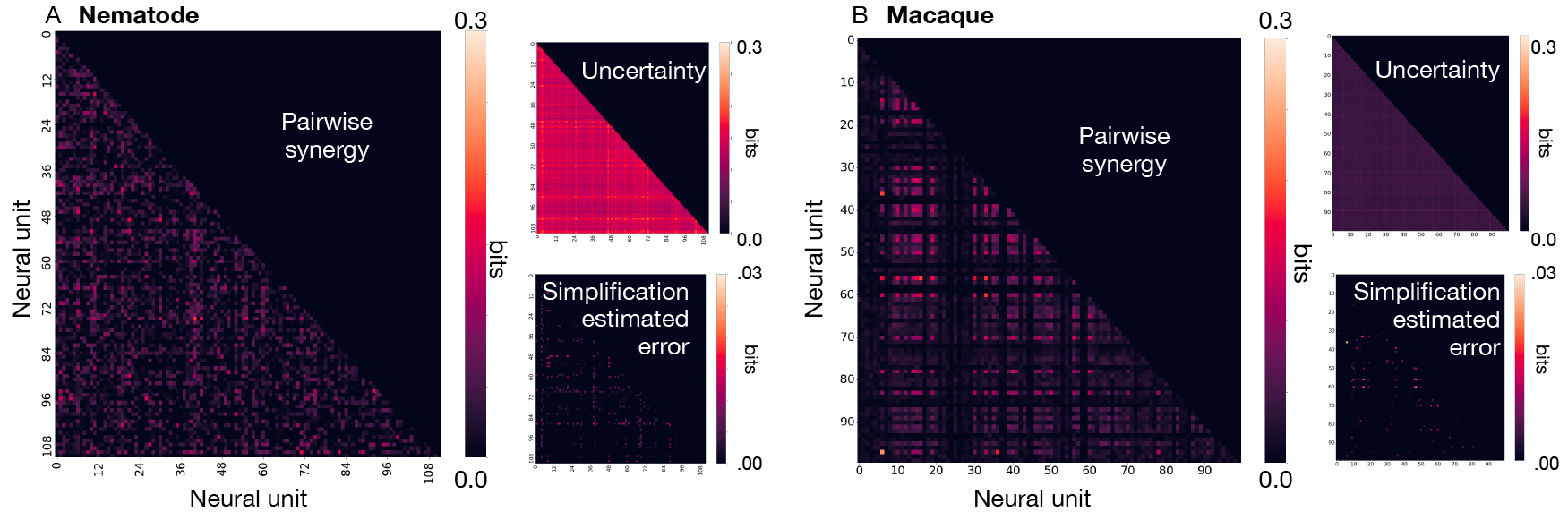
Some pairs of neurons contain synergistic information that is not available at the level of individual neurons. In the macaque just after the decision behavior (B), a number of neuron pairs provide ∼ 0.2 bits of synergistic information. This is estimated using our simplified version of the partial information decomposition. The estimated uncertainty of the total joint entropy, shown in “Uncertainty” subplots, dominates any pairwise synergy in the nematode (A), but is relatively small in the macaque where we have more data. A rough estimate of the error due to using a simplified calculation of synergy, shown in “Simplification estimated error” subplots, is an order of magnitude smaller (note different scale bar; see Methods for details).

To explore how these pairs of neurons encode synergistic information, we plot the pairwise activity states in two dimensions, with colors representing the discrete behavioral decision states (Fig. 3). Our definition of synergy depends on the existence of neural states that, when measured simultaneously, are informative about behavior much more than the respective states measured in each neural unit individually. We highlight an example neural state in each case that contributes to the synergy (orange stars). In the most synergistic pair in the macaque example (Fig. 3D), the geometry of neural states corresponding to each behavior are relatively simply characterized, with two clouds of states separated roughly by a diagonal line. States near the middle of this diagonal contribute to the synergy: These states are more ambiguous at the individual level, with marginals that overlap in both single-unit directions, while knowing the state of both neural units simultaneously is better able to resolve on which side of the line is the joint activity.

**Figure 3.**
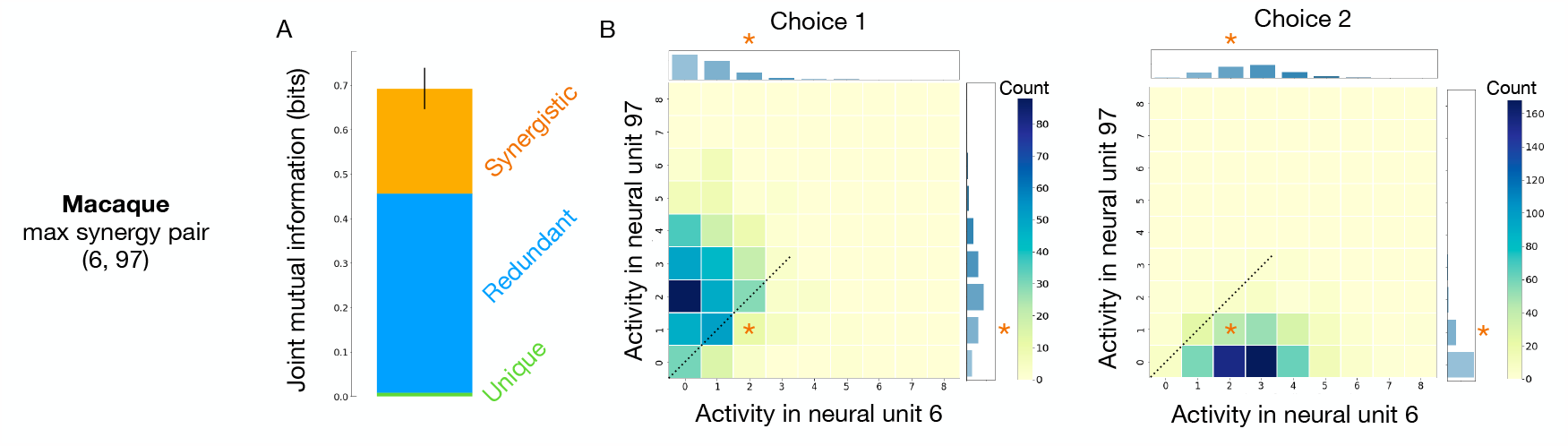
Characterizing the encoding of synergy in a pair of neurons. (A): In the macaque, the partial information decomposition for the joint behavioral information carried by the pair of neurons with maximal synergy. Black vertical errorbar indicates the estimated uncertainty in the total joint entropy. (B): For the same pairs of neural units with largest synergy, we compare the neural state space that is occupied in different behavioral states (two opposite saccade choices) for a visual representation of how that synergy is encoded. Orange stars indicate key neural states that demonstrate synergy. These highlighted neural states are more informative about the behavior when both neurons are measured simultaneously (with large differences across the two main 2D histograms) than when the neurons are measured individually (with small differences across each of the two subplot 1D histograms that represent marginals for each neuron individually). The diagonal dotted lines show a rough delineation between neural activity in the two behavioral choice states.

### 3.3 Information in larger collections of neurons: Linear discriminant analysis (LDA)

Moving finally to arbitrarily large subgroups of neural units, we measure how well we are able to predict behavioral decisions by combining neural units that are individually more or less informative.

In Fig. 4, we plot the performance of LDA in predicting nematode behavior as we change the number of included neurons, for one choice of training and test data. Using activities of the five neurons that are individually most informative (those highlighted in Fig. 1A) produces an accuracy of 0.90 (horizontal red line). At the other extreme, using the activities of all 112 neurons increases the accuracy only slightly to 0.93 (horizontal green line). If we instead choose neurons randomly, increasing the number of neurons gradually improves performance (orange curve). Finally, to better highlight information that is available only at larger scales, we choose neurons randomly while excluding the five most informative ones (blue curve). The intersection of the blue and red curves shows that equal predictive power can be obtained by selecting either five specific highly informative neurons or about 65 random neurons that are individually less informative. For other choices of training and test data, the crossover point varies (Fig. 7), but there is always some number of less informative neurons that performs as well as the five most informative ones.

**Figure 4.**
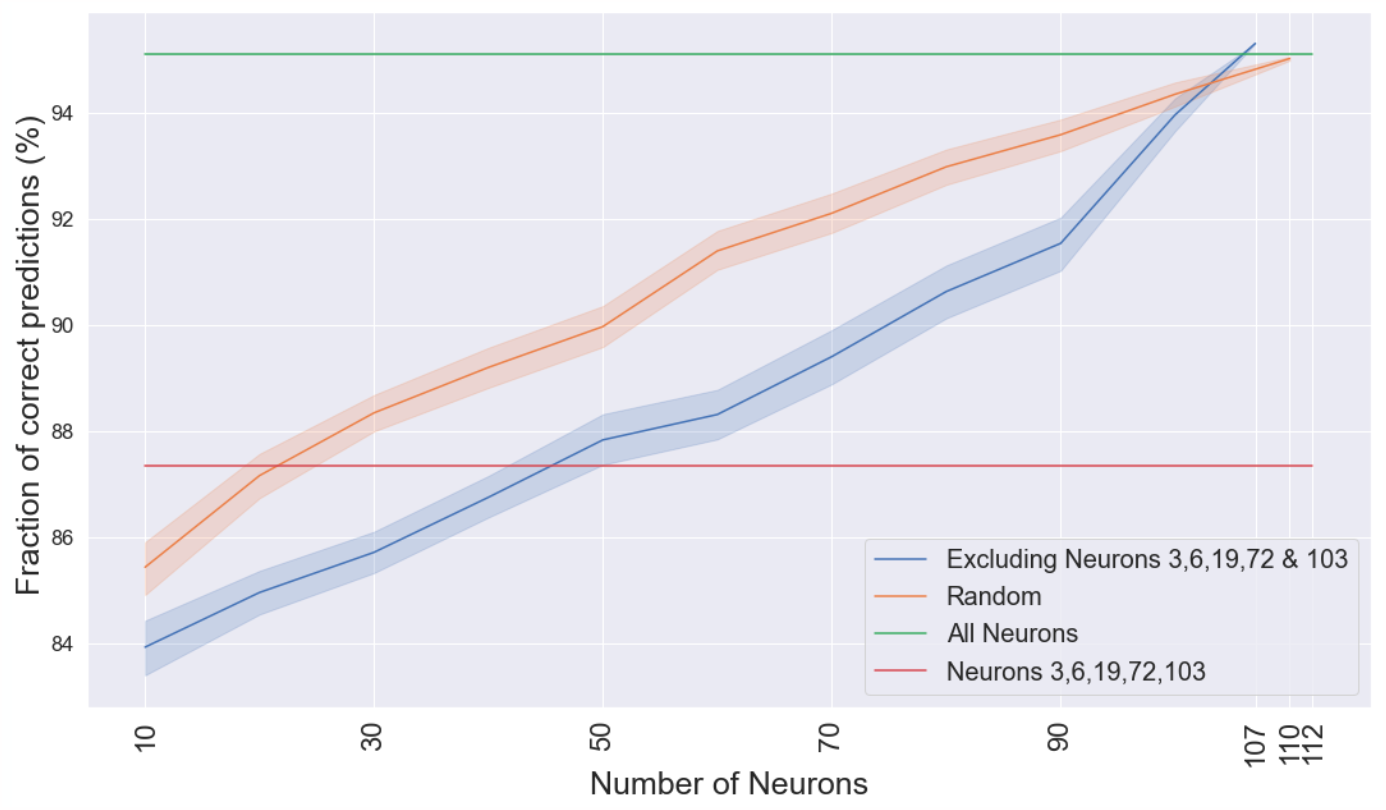
In the nematode, a collection of many individually uninformative neurons is as predictive as a small number of highly informative neurons. The mean accuracy of LDA predictions for forward versus non-forward behavior increases with the number of sampled neurons. Here we compare randomly selected sub-populations of neurons, chosen either from all measured neurons (orange) or from all measured neurons excluding the five that are individually most informative (blue). Measuring more than about 65 individually uninformative neurons (blue) is on average more predictive than the five neurons that are individually most informative (horizontal red line; neurons highlighted in Fig. 1A). Shaded regions indicate ± 1 standard deviation of the mean over 200 choices of random neurons. See Fig. 7 for the same plot across multiple choices of test and training subsets.

In the macaque example data we have an additional dimension of time: neurons are measured both before and after the go cue that directs the macaque to output its decision. In Fig. 5, we show how localized and distributed information coexist as a function of trial time. Near the time of the decision action (about 250 ms after the go cue), the joint activity of the five most informative neurons (red curve) has predictive power that is similar to 50 randomly chosen neural units that exclude those five highly informative units (blue curve). Looking across time, the dominance of localized and distributed information switches: Before the action, the five most informative units are more predictive, and after the action, the combination of individually uninformative units are more predictive.

**Figure 5.**
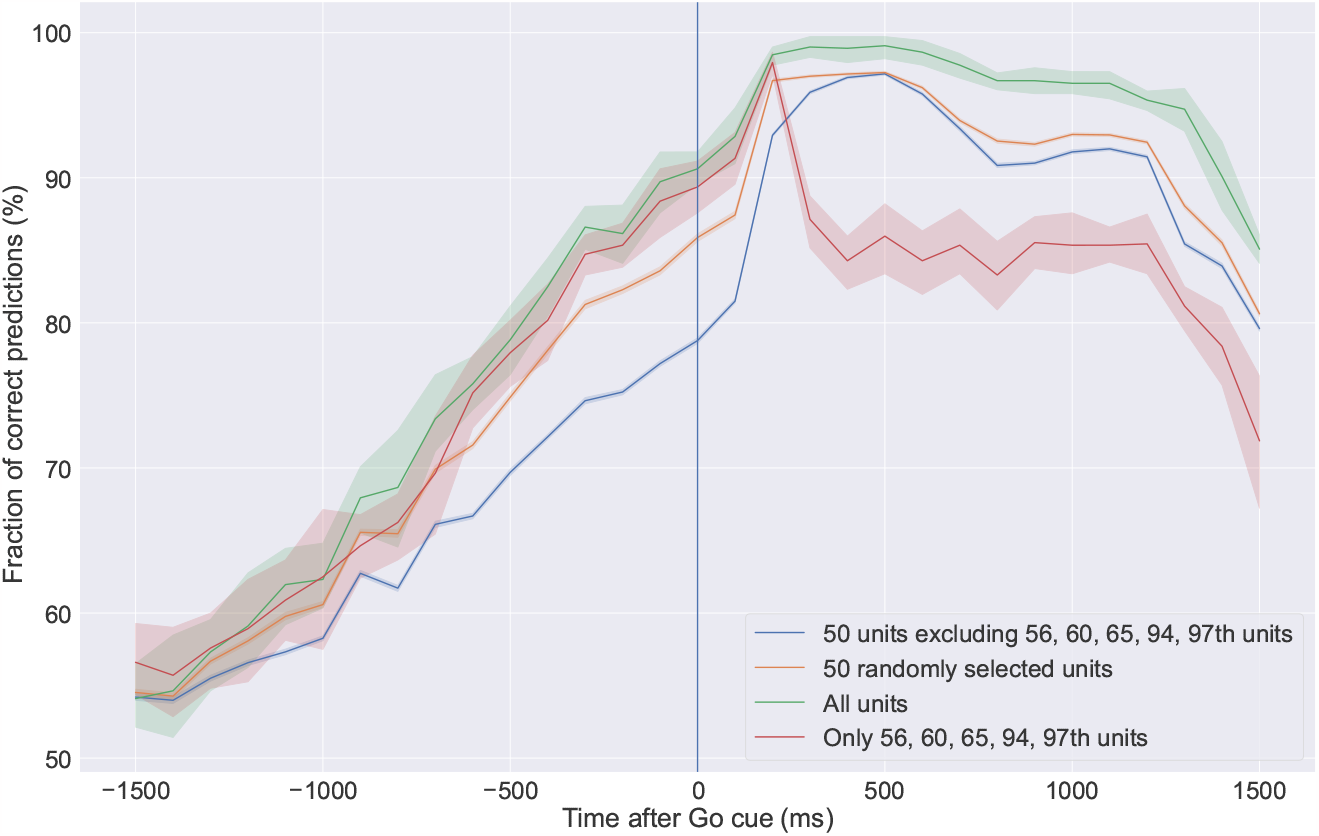
In the macaque, the localization of decision information undergoes a dynamic switch. The ability of LDA to predict behavioral choices as a function of trial time shows that five highly informative neural units (red curve; neural units highlighted in Fig. 1C) contain the most predictive information before the decision action. This switches near the time of the action (about 250 ms after the go cue), when 50 neurons chosen randomly from those that are individually less informative (blue line) are collectively more predictive. Shaded regions indicate ± 1 standard deviation of the mean over 200 choices of random neurons times 4 choices of randomly selected test and training subsets. See Fig. 8 for the same plot across multiple choices of test and training subsets.

## 4 Discussion

Where behavioral information can be found in the brain is a step toward understanding mechanisms of collective decisions—how dynamics of the neural substrate lead to symmetry breaking at the aggregate scale.

We have shown one way to quantify behavioral information across scales, from the level of individual neurons to pairs to larger groups. Our results show that significant amounts of information about behavior can be found redundantly across many scales.

That is, individual neural units can predict behavior (as measured by mutual information); in some cases, pairs of units can predict behavior, even among those that are less predictive individually (as measured by pairwise synergy); and larger subsets of units can predict behavior, even among those that are less predictive individually (as measured by a linear classifier).

In both the nematode and macaque example cases that we study, we find localized behavioral information: some individual neurons carry a significant amount of information. In some cases, behavioral information is both localized and distributed. Specifically, when measured at the same time as the behavioral output, large sets of neurons that exclude the highly informative ones (around 40 to 100 neurons, chosen randomly in the nematode case and from an informative subset in the macaque case) are able to provide as much accuracy as measuring the neurons with the most information. In the macaque example, due to a different experimental design, we are additionally able to characterize information as a function of time relative to the decision. Across all relative times, we find that localized and distributed information coexist, but their relative magnitudes change: A smaller set of specific neurons predict better before the decision action, and large sets of randomly chosen neurons predict better after. Overall, in terms of determining neural encodings and controllers of behavior, our results emphasize that finding behavioral information that is localized does not rule out the possibility of coexisting distributed information.

Such a view is emphasized in population models of neural dynamics in which the stable dynamical states of the system as a whole are foregrounded instead of a design that focuses on the contributions of individual neurons [37, 38]. It is known across many examples in neuroscience that even randomly selected neurons (often within a known brain area, but otherwise not carefully selected) often contain predictive information about behavior. When including enough neurons, typically tens or hundreds, population activity can be used to make good predictions about behavior, in primates [39], leeches [40], and mice [41].

To better characterize the encoding of behavioral information at larger scales, we measured the degree to which the information was synergistic (non-decomposable). In the macaque example, we find significant synergy at the level of pairs of neural units, with some pairs carrying an amount of purely synergistic information (0.24 ± 0.05 bits) that is comparable in magnitude to the most informative individual units (0.45 ± 0.03 bits). For larger groups of neurons, we did not try to measure the synergy because uncertainty in the values of individual mutual information is large enough that this becomes difficult: for a group of 50 neurons that collectively encode 1 bit of information, we would need to be able to measure the information carried by individual units with a resolution of roughly 1/50 = 0.02 bits to start to distinguish unique information from synergistic. While synergy is difficult to define at these larger scales, our results make clear that groups of neurons that individually carry little information and would seem to be lost in the noise can still become highly informative as a group. This is consistent with other work that obtains a rough measure of synergy in larger groups using the O-information, which finds in human fMRI data [24] and macaque single-cell data [25] that larger groups can be found that collectively encode more information compared to the individual neurons or neural regions that constitute the groups.

The dynamics of information localization hints at the mechanisms at work. The collective algorithms that carry out decisions may involve localized information being amplified to many other neurons (as seems to be the case in the macaque example), information coexisting at multiple scales (as seems to be the case in the nematode example), or distributed information that becomes consolidated in localized populations. Going beyond descriptions of information flow, as we measure here, to causal mechanisms will involve characterizing the localization of decision control. Note that having predictive information about a decision is a necessary but not sufficient condition for having control over that decision: The localization of information puts an upper bound on the localization of control. As we sketch in Fig. 6, different mechanisms of decision-making correspond to variation in the sizes of neural populations that have control over the decision (represented by shades of blue). It is possible that the planning and execution of behavioral outputs could be controlled by a sequence of individual neurons, perhaps representing increasing commitment to a given decision along a series of specific neurons [6] (top row). In contrast, a “vote aggregation” mechanism is dependent on the state of a large number of neurons that is then consolidated into a single neuron that stimulates the behavioral action. It is also possible that control remains highly distributed throughout the decision process, with sensory areas less well distinguished from motor areas (as might be suggested by the recent focus on “mixed selectivity” [42]) (bottom row). These cases bound many other graded possibilities.

**Figure 6.**
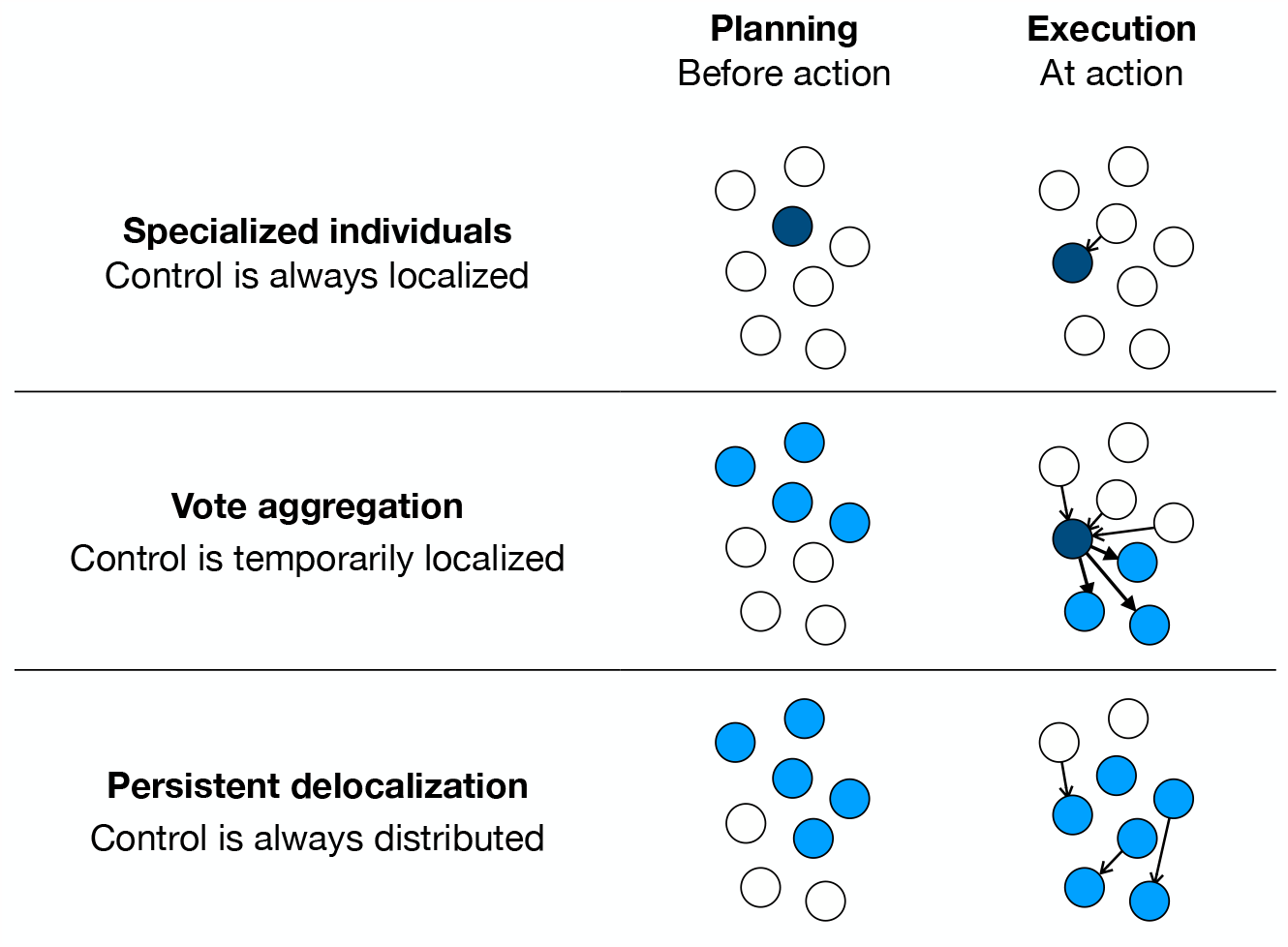
Characterizing mechanisms for distributed decision-making based on localization of control. Circles represent neurons or neural populations, and shades of blue represent control over the eventual decision — light blue distributed, and dark blue localized.

The localization of control has practical implications for how distributed computations work. In particular, theory predicts that timescales of information integration depend on the number of neurons over which that information accumulates [43, 44]. Relatedly, the dynamics of the localization of information is thought to be connected to the motion of the system with respect to collective transitions [5]. Collective computation in social systems can also be implemented by multiple dynamical mechanisms for coming to a consensus decision, and these mechanisms differ in the dynamics of the informational content of the states of individual components [45, 46].

Techniques in control theory often search for a specific, small set of components that can optimally control collective outcomes. Such control can be studied with respect to coarse-grained collective decision states as we study here (e.g. [47, 48]) or more fine-grained behavioral quantities down to the level of all possible behaviors allowed by the motor system (e.g. [8, 49]). Across such examples, an implicit assumption is that finding smaller sets of controlling components is desirable because sparse control is easier to implement. Yet the existence of information about behavioral outputs distributed among larger subsets, even when chosen randomly among components that individually contain little information, suggests the possibility of alternative control strategies. In both example systems that we study here, our linear encoding (via LDA) successfully captures most of the categorical behavioral information when including ∼ 50 neurons, suggesting that nonlinear methods for decoding may be unnecessary when including enough individual neurons. This is reminiscent of Cover’s theorem in machine learning, in which linear classifiers can become very accurate when the system’s state is transformed into a higher-dimensional space [50]. If control could equally well be implemented via a linear perturbation of a larger set of randomly selected neurons, this would eliminate the need for a potentially costly optimization search over a combinatorially large space of control subsets and associated nonlinear perturbations. That is, it is possible that as control strategies become more distributed, they are more difficult in the sense of requiring inputs to more neurons but less difficult in the sense of being easier to find.

Causal network inference is particularly difficult in collective systems due to the curse of dimensionality, and producing an inferred network that makes reasonable predictions typically relies on simplifying assumptions, sometimes under the guise of statistical priors or regularizers [51, 52]. Often control is attributed to the smallest number of components possible, as this greatly simplifies inference. On the other hand, of fundamental interest to complex systems science are cases in which causal determinants are distributed over many components [22,53–56]. When using inference methods with a bias toward sparsity of individual components, it is possible that distributed mechanisms could be overlooked. As one alternative, sparse methods can be designed to search for explanations in terms of a small number of basis functions that each involve an arbitrary number of components [57].

The “selectivity” of individual neurons, with activity that is informative of particular sensory or behavioral states, may be a property even of randomly wired networks [58]. Such results shift the discussion from the properties of precise networks that produce adaptive dynamics in a predefined way (and often involving particular neurons or groups of neurons predefined to represent a particular piece of information) to random network ensembles that have fewer constraints on particular structure at the micro scale but can reliably perform a task at the macro scale [59]. This might correspond to cases like ours in which information can exist both highly concentrated in some neurons and widely dispersed in others, in that randomly wired networks are perhaps more likely to contain many coexisting (redundant) encodings at a variety of scales, whereas engineered systems are more likely to encode information at the individual component scale because this makes them easier to understand.

In sum, our perspective challenges the implicit narrative that we are looking for the single “correct” scale at which to analyze the neural systems that control behavioral decisions. Instead, information, and perhaps control, can exist simultaneously at multiple scales.

## Supplementary Figures

**Figure 7.**
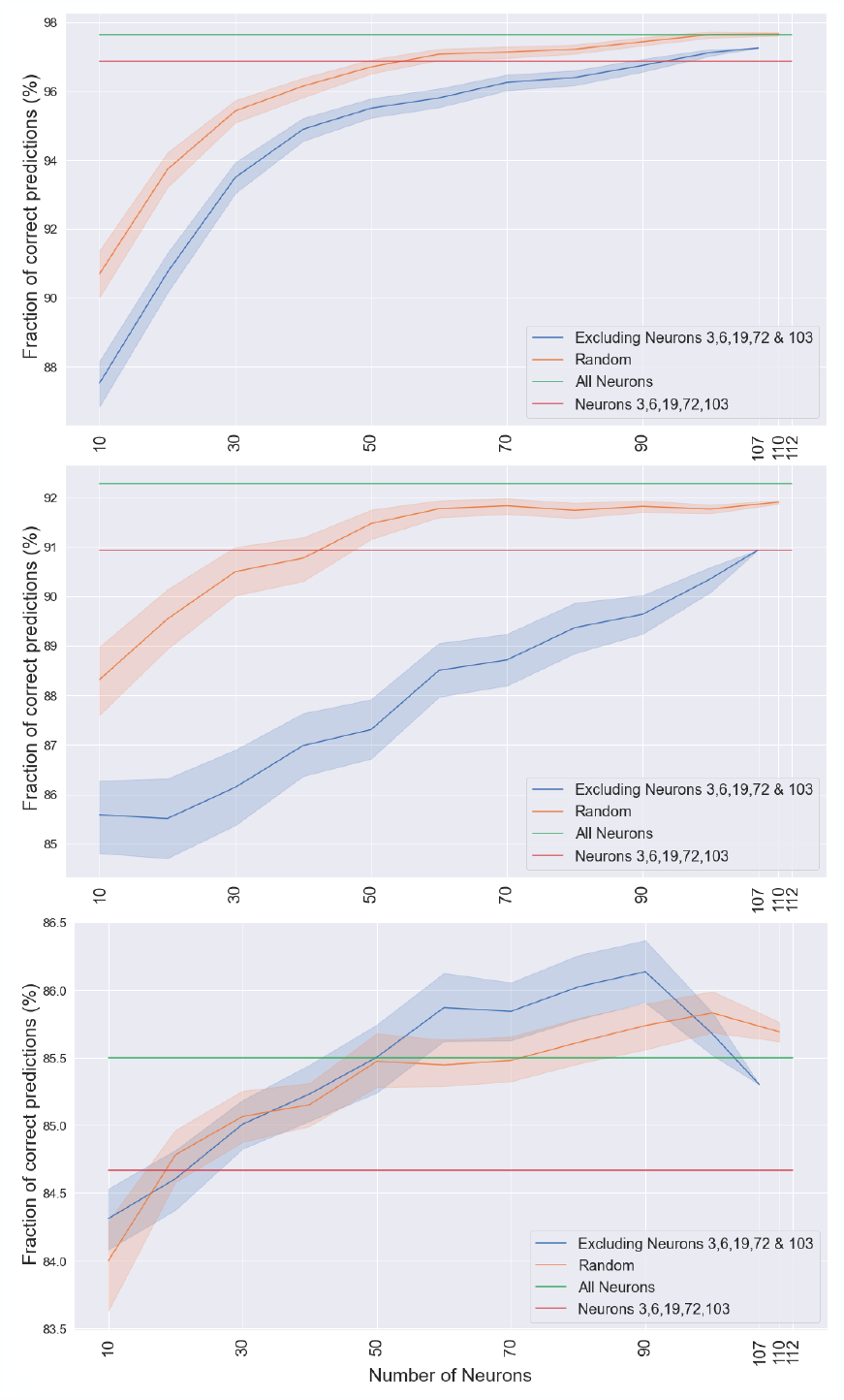
The degree of predictive power depends on which portion of the nematode time series is held out for testing. Remaking Fig. 4 for three other splits of the nematode time series into training and test data, we see that the overall degree of predictability is highly dependent on this choice. Still, our qualitative conclusion holds: In each case, randomly chosen less-informative neurons (blue curve) are able to match the predictive power of the five most informative neurons (red line) when enough of them are included.

**Figure 8.**
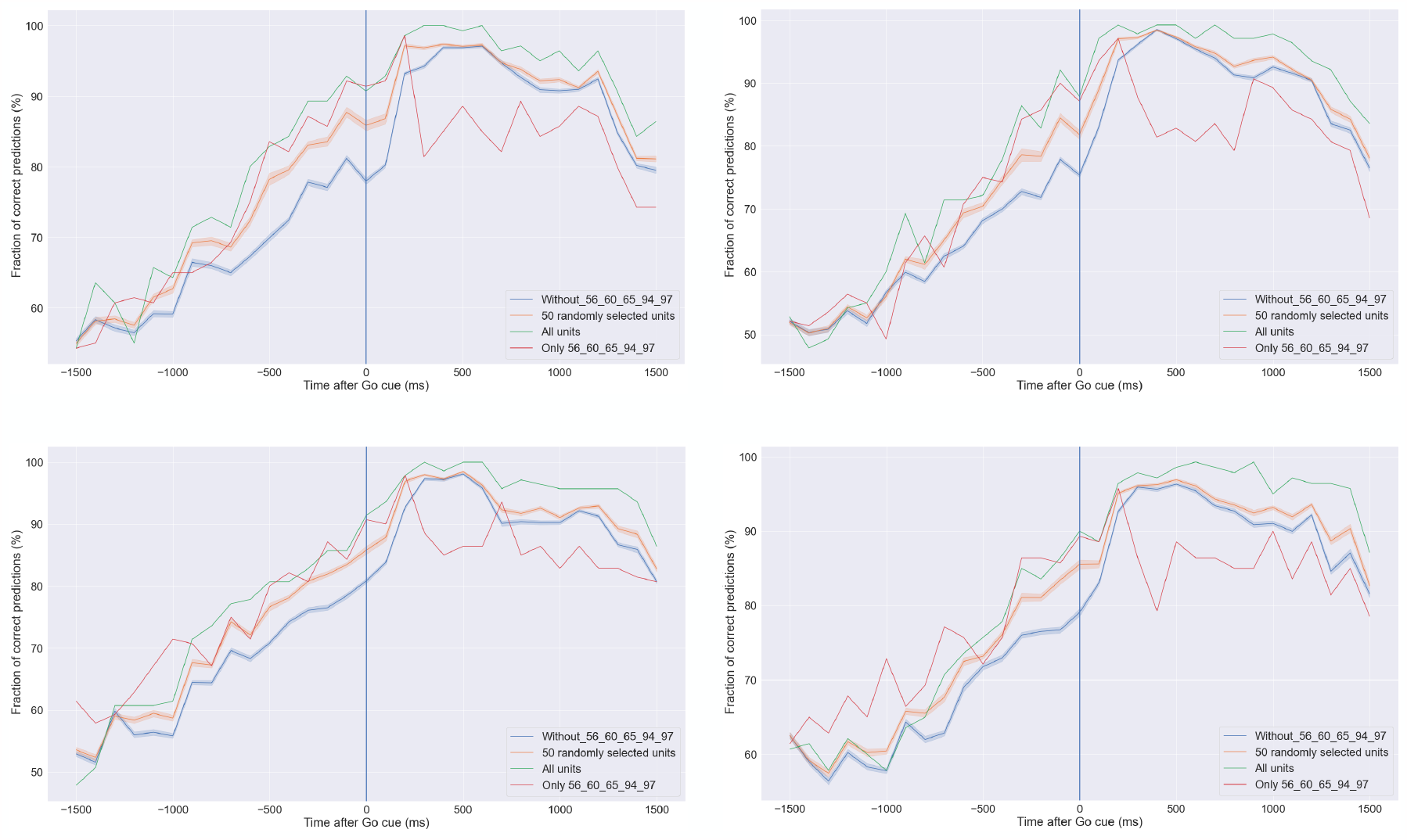
The degree of variation caused by the choice of training and test data in the macaque case for neurons with the largest information averaged over trial time. Here we show three versions of Fig. 5 for three separate splits of the macaque data into training and test sets.

**Figure 9.**
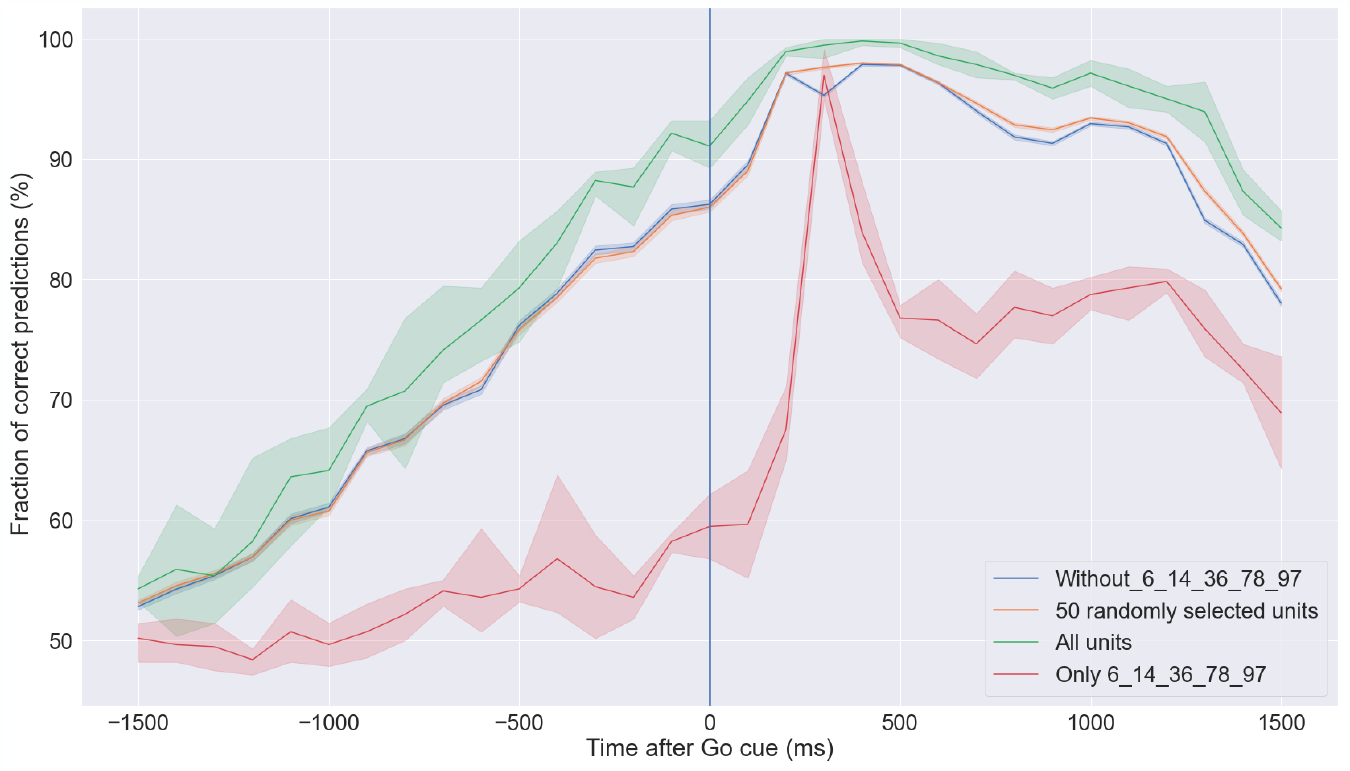
In the macaque case, some neurons are highly informative only near the time of the output behavior. Here we remake Fig. 5 and instead highlight in red the predictive power of five individual neural units with largest mutual information near the time of the action (300 ms post go cue; neural units highlighted in Fig. 1B), as opposed to selecting those with the largest mutual information averaged over trial time. In this case, the selected neural units are as predictive as other subsets only near that single time.

**Figure 10.**
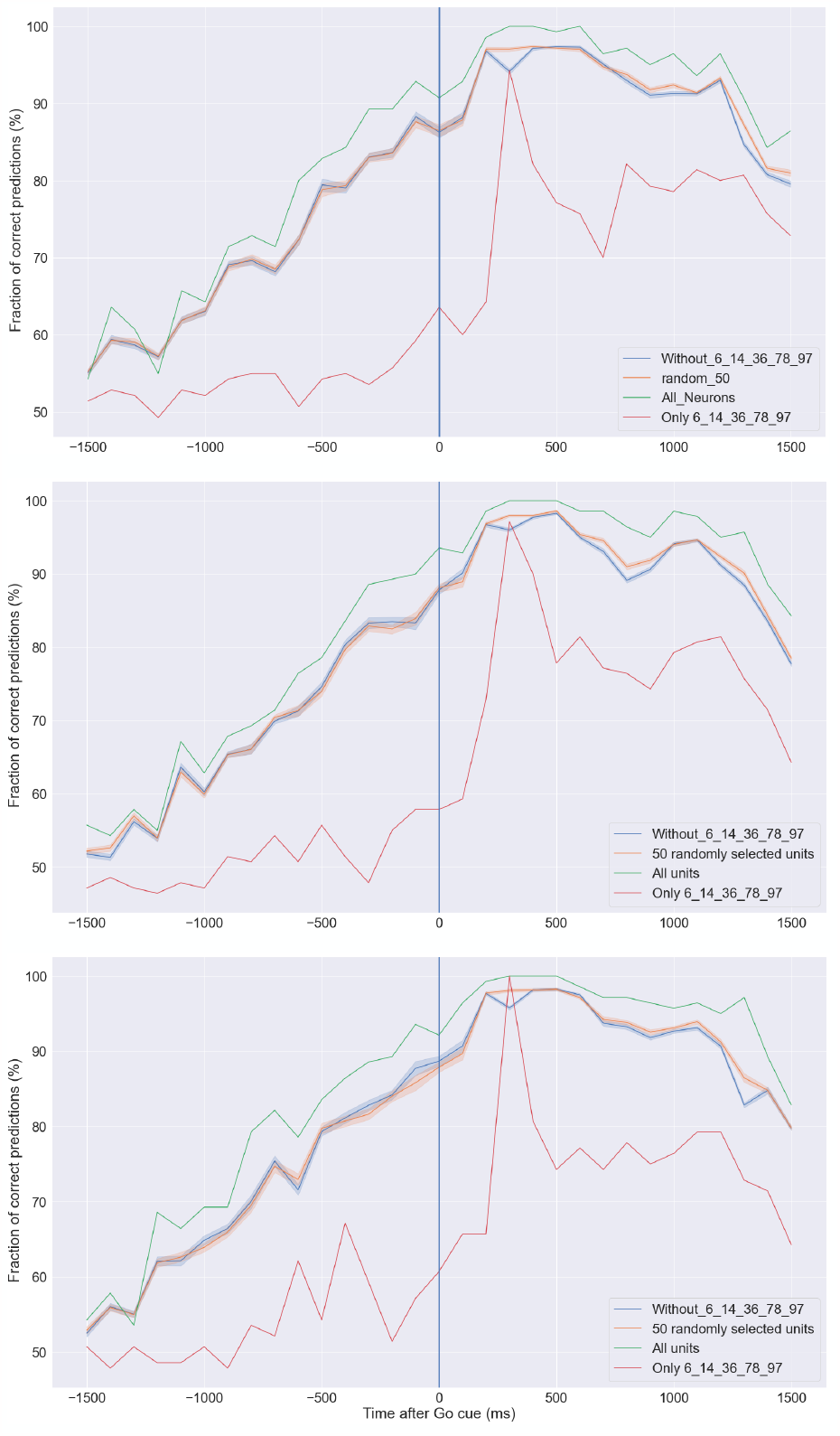
The degree of variation caused by the choice of training and test data in the macaque case for neurons selected near the time of the action. Here we show three versions of Fig. 9 for three separate splits of the macaque data into training and test sets.

